# Enhancers facilitate the birth of *de novo* genes and their integration into regulatory networks

**DOI:** 10.1101/616581

**Authors:** Paco Majic, Joshua L. Payne

## Abstract

Regulatory networks control the spatiotemporal gene expression patterns that give rise to and define the individual cell types of multicellular organisms. In eumetazoa, distal regulatory elements called enhancers play a key role in determining the structure of such networks, particularly the wiring diagram of “who regulates whom.” Mutations that affect enhancer activity can therefore rewire regulatory networks, potentially causing changes in gene expression that are adaptive. Here, we use whole-tissue and single-cell transcriptomic and chromatin accessibility data from mouse to show that enhancers play an additional role in the evolution of regulatory networks: They facilitate network growth by creating transcriptionally active regions of open chromatin that are conducive to *de novo* gene evolution. Specifically, our comparative transcriptomic analysis with three other mammalian species shows that young, mouse-specific intergenic open reading frames are preferentially located near enhancers, whereas older open reading frames are not. Mouse-specific intergenic open reading frames that are proximal to enhancers are more highly and stably transcribed than those that are not proximal to enhancers or promoters, and they are transcribed in a limited diversity of cellular contexts. Furthermore, we report several instances of mouse-specific intergenic open reading frames that are proximal to promoters that show evidence of being repurposed enhancers. We also show that open reading frames gradually acquire specific interactions with enhancers over macro-evolutionary timescales, helping integrate new genes into existing regulatory networks. Taken together, our results highlight a dual role of enhancers in expanding and rewiring gene regulatory networks.

## Introduction

Enhancers are a defining characteristic of eumetazoan gene regulatory networks. They recruit transcription factors and cofactors that “loop out” DNA to bind core promoters and increase the expression of target genes (Catarino and Stark 2018; Haberle and Stark 2018), thus mediating interactions between genes. Such interactions are highly dynamic throughout development, facilitating the differential deployment of distinct regulatory sub-networks in different cells, which helps define cell-type specific spatiotemporal gene expression patterns (Davidson and Levine 2008; Spitz and Furlong 2012).

Enhancer activity is not only dynamic throughout development, but also throughout evolution (Villar, et al. 2015). The reason is that mutations in enhancer sequences can create or ablate interactions with regulatory proteins, thus enabling modifications in gene use without affecting gene product (Prud’homme, et al. 2007; Carroll 2008). Such changes alter a regulatory network’s wiring diagram of “who regulates whom,” which can cause changes in gene expression patterns that embody or lead to evolutionary adaptations or innovations (Peter and Davidson 2011). Examples include the archetypical pentadactyl limb anatomy of extant tetrapods (Kherdjemil, et al. 2016), ocular regression in subterranean rodents (Partha, et al. 2017; Roscito, et al. 2018), limb loss in snakes (Kvon, et al. 2016; Roscito, et al. 2018), convergent pigmentation patterns in East African cichlids (Kratochwil, et al. 2018), the diversity of butterfly wing patterns (Barton, et al. 2016), the mammalian neocortex (Emera, et al. 2016), and cell type diversity in eumetazoans (Sebé-Pedrós, et al. 2018).

Regulatory networks not only evolve via rewiring, but also via the addition of new genes (Teichmann and Babu 2004). Gene duplication, retrotransposition, gene fusion, the domestication of genomic parasites, and horizontal gene transfer are all means by which new genes can arise from pre-existing genes (Kaessmann 2010), and thus expand gene regulatory networks. In addition, it is becoming increasingly appreciated that new genes can arise *de novo* from non-coding regions of the genome (Carvunis, et al. 2012; Betran, et al. 2013; Li, et al. 2014; Kim, et al. 2015; McLysaght and Hurst 2016; Van Oss and Carvunis 2019; Willemsen, et al. 2019). For protein-coding genes, the essential prerequisites of this process are the formation of an open reading frame (ORF), together with the transcription and translation of that ORF. Because much of the genome is transcribed (Kapranov, et al. 2007; Neme and Tautz 2016) and many lineage-specific transcripts containing ORFs show evidence of translation (Wilson and Masel 2011; Ingolia, et al. 2014; Ruiz-Orera, et al. 2014; Prabh and Rödelsperger 2016; Ruiz-Orera, et al. 2018; Schmitz, et al. 2018; Ruiz-Orera and Alba 2019; Zhang, et al. 2019), the *de novo* evolution of new protein-coding genes is also a likely contributor to the growth of gene regulatory networks.

An important question concerning *de novo* genes is how they integrate into existing regulatory networks, and what role enhancers may play in this process. It has been hypothesized that enhancer acquisition allows new genes to expand their breadth of expression, providing opportunities to acquire new functions in different cellular contexts (Tautz and Domazet-Loso 2011). Enhancers may therefore help new genes integrate into existing regulatory networks via edge formation and rewiring. Less appreciated is the role enhancers may play in the origin of *de novo* genes (Wu and Sharp 2013), and thus in the growth of gene regulatory networks. The physical proximity between active enhancers and their target genes (Levine, et al. 2014) – facilitated by DNA looping – creates a transcriptionally permissive environment that is engaged with RNA polymerase II, which may lead to the transcription of DNA near the enhancer, or to the transcription of the enhancer itself, producing so-called enhancer RNA (De Santa, et al. 2010; Kim, et al. 2010; Li, et al. 2016; Haberle and Stark 2018). If the transcript contains an open reading frame, then such increased transcription will increase the likelihood of interaction with ribosomes, and because enhancers are typically active in a small number of cell types, interactions with ribosomes will occur in a limited diversity of cellular contexts. This may help purge toxic peptides and enrich for benign peptides, a process that has been hypothesized to increase the likelihood of *de novo* gene birth (Wilson and Masel 2011). Moreover, similarities in the architectures of enhancers and promoters may facilitate the regulatory repurposing of the former into the latter (Carelli, et al. 2018), reinforcing the transcription of new open reading frames that emerge near enhancers. Thus, enhancers may play a dual role in the evolution of *de novo* genes, and consequently in the evolution of gene regulatory networks. By creating a transcriptionally permissive environment, enhancers may facilitate the origin of *de novo* genes; by physically interacting with gene promoters, enhancers may facilitate the integration of *de novo* genes into existing regulatory networks.

The first evidence that enhancers can facilitate *de novo* gene birth was recently provided using whole-animal transcriptomic and epigenetic data from the nematode *Pristionchus pacificus* (Werner, et al. 2018). Specifically, the transcription start sites of expressed genes that were in open chromatin and private to *P. pacificus* were found to be in closer proximity to histone modifications indicative of enhancers than the transcription start sites of expressed genes that were in open chromatin and shared with other nematode species. While this evidence is compelling, additional systematic analyses are required to draw firm conclusions and to address remaining open questions. For example, we do not yet know about the generality of this mechanism, specifically whether it applies to other clades of eumetazoa. Furthermore, information on the stability of the transcribed ORFs or their potential for translation is still lacking. We also do not know about the cell-type specificity of the enhancers that facilitate *de novo* gene birth (because the data used to study *P. pacificus* were derived from the whole animal) or how the facilitating role of enhancers in *de novo* gene birth differs from that of other means of pervasive transcription (Neme and Tautz 2016). Finally, we do not know how enhancers integrate new genes into existing cellular networks, especially over macro-evolutionary timescales.

Here, we take an integrative approach to address these open questions and to study the potential dual role of enhancers in the evolution of gene regulatory networks. We leverage whole-tissue and single-cell transcriptomic and functional genomics data from mouse that describe gene expression levels, chromatin accessibility, and chemical modifications to histones, as well as phylostratigraphic estimates of the ages of transcribed ORFs. We find young ORFs are preferentially located near enhancers, whereas older ORFs are not. Some of these young ORFs likely are enhancers, as evidenced by their bidirectional transcription – a hallmark of enhancer activity. Mouse-specific intergenic ORFs that are proximal to enhancers are more highly and stably transcribed than mouse-specific intergenic ORFs that are not proximal to enhancers or promoters, and they are transcribed in a limited diversity of cellular contexts, thus highlighting fundamental differences between the facilitating role of enhancers versus other forms of pervasive transcription in *de novo* gene birth. We find the transcripts of enhancer-proximal ORFs often associate with ribosomes, and we uncover several instances of mouse-specific intergenic ORFs that are proximal to promoters that are likely repurposed enhancers. Finally, we show the number of enhancer interactions per ORF increases with ORF age, which correlates with an increase in expression breadth, even across macro-evolutionary timescales. In sum, our findings support a dual role for enhancers in the origin of *de novo* genes and in their integration into gene regulatory networks.

## Results

### Mouse-specific intergenic ORFs are often proximal to enhancers

We considered a set of 56,262 ORFs from transcripts expressed in the liver, brain, and testis of *Mus musculus.* Previous work assigned phylogenetic ages to these ORFs (Schmitz, et al. 2018), based on the presence of homologous sequences in the transcriptomes of other mammalian species, including rat, human, and opossum (Fig. 1A). We further classified the mouse-specific ORFs as genic or intergenic, based on whether or not they are proximal to older, annotated genes (Materials and Methods). We use the term *proximal* to mean within 500bp (in the supplementary material, we show our findings are qualitatively insensitive to changing this definition to 250bp and 1000bp, see Figs. S4–S8), and we use an ORF’s first exon to calculate its distance from other genomic features. To characterize the regulatory background of an ORF, we considered data describing histone modifications that are indicative of promoters and enhancers (Heintzman, et al. 2007). Specifically, we merged chromatin immunoprecipitation followed by DNA sequencing (ChIP-seq) data for H3K27ac, H3K4me1, and H3K4me3 obtained from 23 mouse tissues and cell types (ENCODE 2012). We considered enhancers to be those genomic regions where H3K27ac and/or H3K4me1 peaks do not overlap H3K4me3 peaks in any tissue, and promoters to be those genomic regions with H3K4me3 peaks (Creyghton, et al. 2010; Berthelot, et al. 2018) (Materials and Methods).

**Figure 1.**
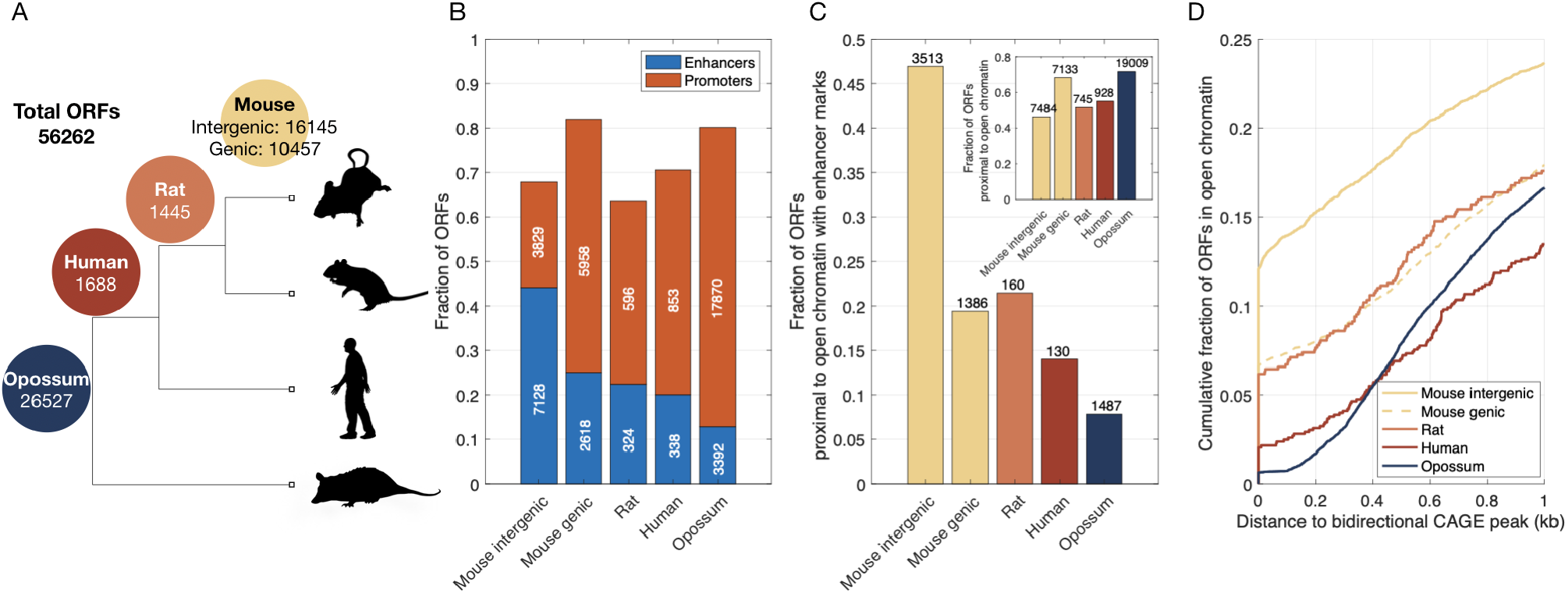
Mouse-specific intergenic ORFs are often proximal to enhancers. A) Phylogeny showing the four age classes of the 56,262 ORFs. The numbers on the branches indicate the number of ORFs that are either mouse-specific or shared with rat, human, and opossum. Mouse-specific ORFs are further classified as intergenic or genic. B) Fraction of ORFs that are proximal to ChIP-seq peaks indicative of enhancers (H3K27ac and/or H3K4me1 without overlapping H3K4me3) or promoters (H3K4me3), shown in relation to ORF class. C) Fraction of ORFs that are proximal to regions of open chromatin that contain enhancers, but not promoters, shown in relation to ORF class. The inset shows the fraction of ORFs that are proximal to regions of open chromatin, regardless of whether those regions contain promoters or enhancers. D) Cumulative fraction of ORFs that are proximal to regions of open chromatin, shown in relation to their distance to the closest bidirectional CAGE peak.

The majority of ORFs in each age class are proximal to a promoter or an enhancer (Fig. 1B). Remarkably, mouse-specific intergenic ORFs are the only class of ORFs that are more likely to be proximal to enhancers than to promoters. While the first exon of nearly 45% (7,128) of mouse-specific intergenic ORFs are proximal to enhancers, fewer than 25% of rat, human, and opossum-shared ORFs are proximal to enhancers. Similar trends are observed when we restrict our attention to ORFs that are within, or proximal to, genomic regions of open chromatin in at least one of thirteen mouse tissues (Fig. 1C; Materials and Methods). Specifically, ∼47% (3,513 out of 7,484) of mouse-specific intergenic ORFs are proximal to regions of open chromatin that harbor histone modifications indicative of enhancers, but not promoters, whereas fewer than 21% of rat, human, and opossum-shared ORFs are proximal to such regions. Similar trends are also observed when we consider histone modification data from individual tissues, as opposed to merging data across cell and tissue types. Specifically, 25% (281 ORFs), ∼36% (897 ORFs), and ∼20% (537 ORFs) of intergenic mouse-specific ORFs that are in open chromatin and expressed in liver, brain, and testis, respectively, are proximal to an enhancer in that tissue, as compared to <10% of genic and older ORFs, which are instead preferentially proximal to promoters (Fig. S2D-F). Finally, mouse-specific intergenic ORFs are more likely to show evidence of bidirectional transcription – a hallmark of enhancer activity (Andersson, et al. 2014) – than any other class of ORFs (Fig. 1D), with 12% of the ORFs overlapping a bidirectional CAGE peak and nearly 20% (1,429) of the ORFs proximal to a bidirectional CAGE peak (Fig. S1 shows that these trends are not driven by exon length). Taken together, these results support a model in which enhancers facilitate the expression of young ORFs (Wu and Sharp 2013; Werner, et al. 2018).

### Intergenic ORFs that are proximal to enhancers are highly and stably transcribed, relative to intergenic ORFs that are not proximal to enhancers or promoters

We next asked what differentiates the facilitating role of enhancers in *de novo* gene birth from other forms of pervasive transcription taking place away from promoters and enhancers. We hypothesized that because enhancers are regularly engaged with the transcriptional machinery, they may confer higher levels of expression and greater expression stability. To test this hypothesis, we compared the expression levels and stabilities of intergenic mouse-specific ORFs that are proximal to enhancers with those of intergenic mouse-specific ORFs that are not proximal to enhancers or promoters, using transcriptomic, histone modification, and chromatin accessibility data from liver, brain, and testis (Materials and Methods).

In all three tissues, we observed that mouse-specific intergenic ORFs that are proximal to enhancers have a higher median expression level than mouse-specific intergenic ORFs that are not proximal to enhancers or promoters (Fig. 2A-C; Wilcoxon signed-rank test, *p* < 0.001 in liver, *p* = 0.003 in brain, *p* = 0.02 in testis). To measure expression stability, we calculated the entropy of expression across biological replicates (Materials and Methods). When this measure equals its minimum of zero, the ORF is expressed in only one of the replicates; when it equals its maximum of one, the ORF is expressed at equal levels across replicates. In all three tissues, we observed that expression stability is higher for mouse-specific intergenic ORFs that are proximal to enhancers than for mouse-specific intergenic ORFs that are not proximal to enhancers or promoters (Fig. 2D-F, Wilcoxon’s signed-rank test, *p* < 0.001). These observations support the hypothesis that enhancers confer higher expression levels and greater expression stability to proximal ORFs than do other forms of pervasive transcription away from promoters and enhancers.

**Figure 2.**
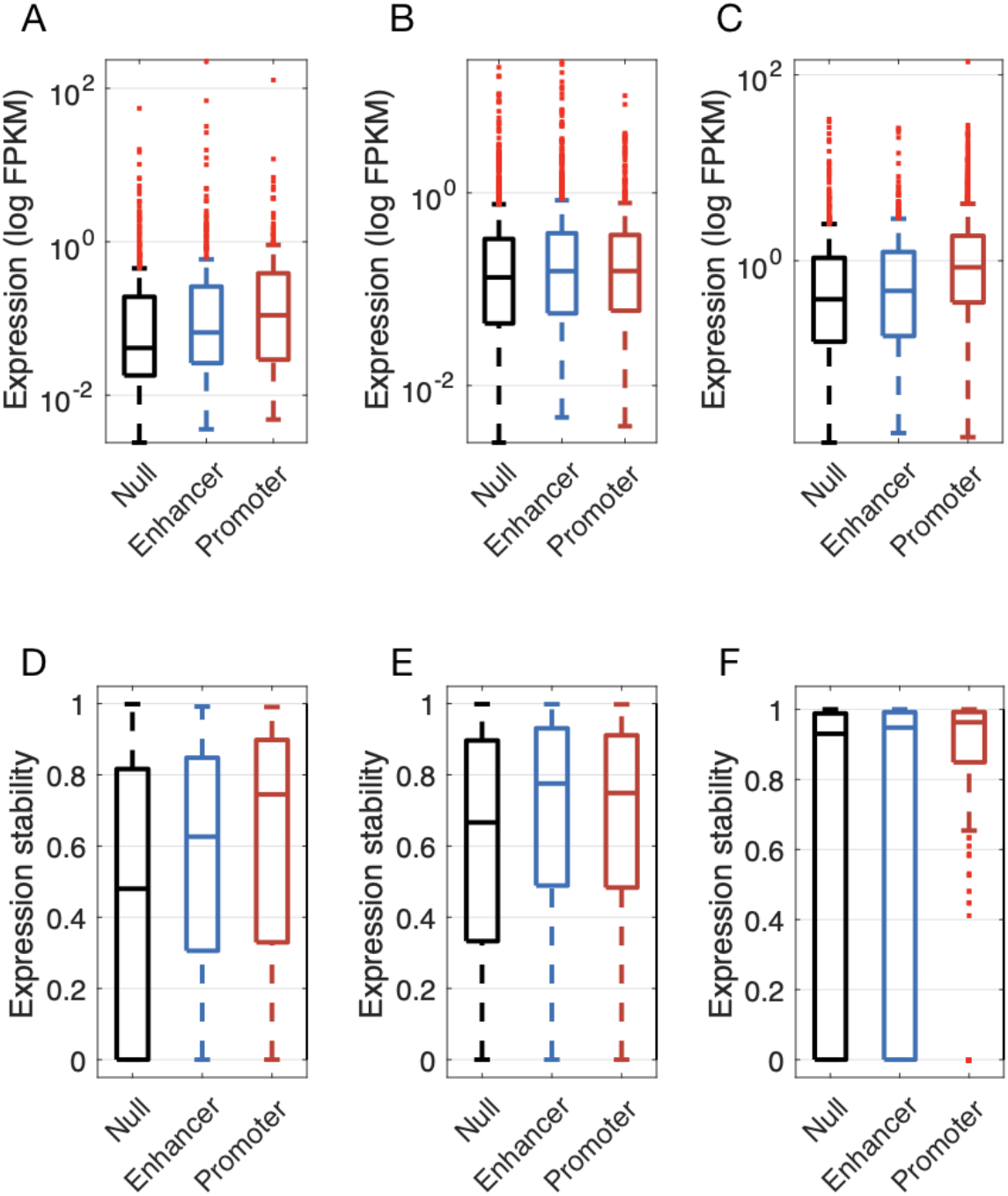
Mouse-specific intergenic ORFs that are proximal to enhancers are more highly expressed and have greater expression stability than mouse-specific intergenic ORFs that are not proximal to enhancers or promoters. Expression level of mouse-specific intergenic ORFs proximal to enhancers, promoters, or neither (‘Null’) in A) liver, B) brain, and C) testis. Expression stability of mouse-specific intergenic ORFs proximal to enhancers, promoters, or neither (‘Null’) in D) liver (8 replicates), E) brain (8 replicates), and F) testis (2 replicates).

### Intergenic ORFs that are proximal to enhancers are more likely to associate with ribosomes than intergenic ORFs that are not proximal to enhancers or promoters

Many non-coding transcripts associate with ribosomes (Wilson and Masel 2011; Ingolia, et al. 2014; Ruiz-Orera, et al. 2014; Zhang, et al. 2015). It has been suggested that this may enrich the pool of transcribed ORFs for benign peptides, thus increasing the likelihood of *de novo* gene birth (Wilson and Masel 2011). We hypothesized that because of their increased expression levels and stabilities, mouse-specific intergenic ORFs that are proximal to enhancers will be more likely to associate with ribosomes than mouse-specific intergenic ORFs that are not proximal to enhancers or promoters. To test this hypothesis, we considered liver, brain, and testis data from a ribosomal profiling assay called ribo-seq, which describes the transcriptome-wide binding patterns of ribosomes to RNA molecules (Ruiz-Orera, et al. 2018; Ruiz-Orera and Alba 2019) (Materials and Methods).

Following Schmitz et al. (2018), we first consider a permissive definition of ribosomal association: at least one read mapping to the first exon of an ORF. We found that mouse-specific intergenic ORFs that are proximal to enhancers are ∼10% more likely to associate with ribosomes than mouse-specific intergenic ORFs that are not proximal to enhancers or promoters, and ∼10% less likely to associate with ribosomes than mouse-specific intergenic ORFs that are proximal to promoters (Fig. 3A). When we apply more conservative thresholds for ribosomal association, mouse-specific intergenic ORFs that are proximal to enhancers remain more likely to associate with ribosomes than mouse-specific intergenic ORFs that are not proximal to enhancers or promoters, and less likely than mouse-specific intergenic ORFs that are proximal to promoters, although the differences in ribosomal association between these classes decreases as the threshold for ribosomal association increases, both when evaluating reads per kilobase mapped to the first exon (Fig. 3A), or simply total number of reads mapped to the first exon (Fig. 3B). These trends remain when considering tissue-specific transcriptomic, histone modification, and ribosomal association data for liver, brain, and testis (Fig. 3C).

**Figure 3.**
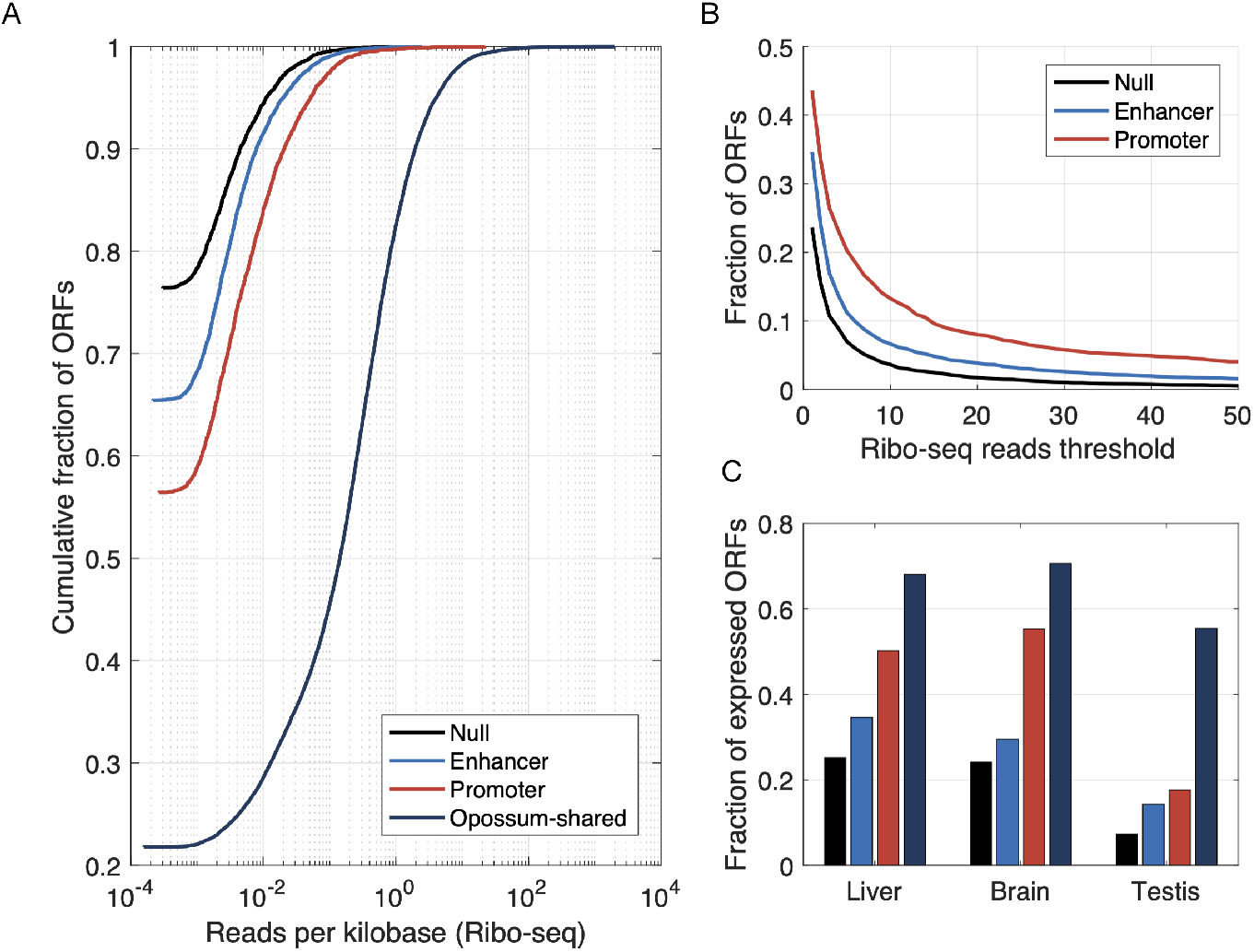
Mouse-specific intergenic ORFs that are proximal to enhancers are more likely to associate with ribosomes than mouse-specific intergenic ORFs that are not proximal to enhancers or promoters. A) Cumulative fractions of mouse-specific intergenic ORFs that are proximal to an enhancer, a promoter, or neither (‘Null’), or to opossum-shared ORFs, shown in relation to the number of ribo-seq reads mapped per kilobase to the first exon of each ORF. B) Fraction of ORFs with ribosomal association, shown in relation to the minimum threshold for the number of reads mapped. C) Fraction of ORFs expressed in liver, brain, and testis for which at least one tissue-specific ribo-seq read could be mapped to their first exon.

### Intergenic ORFs that are proximal to enhancers are expressed in more cellular contexts than intergenic ORFs that are not proximal to enhancers or promoters

In the model of enhancer-facilitated *de novo* gene birth studied here, ORFs emerging near enhancers are likely to have their expression restricted to cells where those enhancers are active. Enhancers are often specific to a small number of cell types (He, et al. 2014), which may reduce the potential for enhancer-proximal ORFs to have deleterious pleiotropic effects, while simultaneously exposing the ORFs to a limited diversity of cellular contexts in which they may confer a selective advantage. To study the breadth of expression of ORFs, we considered two sources of data: whole tissue measurements of total RNA across 10 tissues, and single-cell measurements of open chromatin across 38 cell types (Materials and Methods).

We found that mouse-specific intergenic ORFs that are proximal to enhancers are expressed in more tissues (Wilcoxon’s signed rank test, *p* < 0.001; Fig. 4A) and are in open chromatin in more cell types (Wilcoxon’s signed rank test, *p* < 0.001; Fig. 4B) than mouse-specific intergenic ORFs that are not proximal to enhancers or promoters. However, these ORFs are expressed in fewer tissues (Wilcoxon’s signed rank test, *p* < 0.001; Fig. 4A) and are in open chromatin in fewer cell types (Wilcoxon’s signed rank test, *p* < 0.001; Fig. 4B) than mouse-specific intergenic ORFs that are proximal to promoters, genic ORFs, and ORFs shared with rat, human, and opossum. ORFs emerging near enhancers are therefore transcribed in a limited number of cellular contexts, providing opportunities to purge toxic peptides, and potentially balancing the risk of deleterious pleiotropic effects with the reward of sampling a diversity of cellular environments.

**Figure 4.**
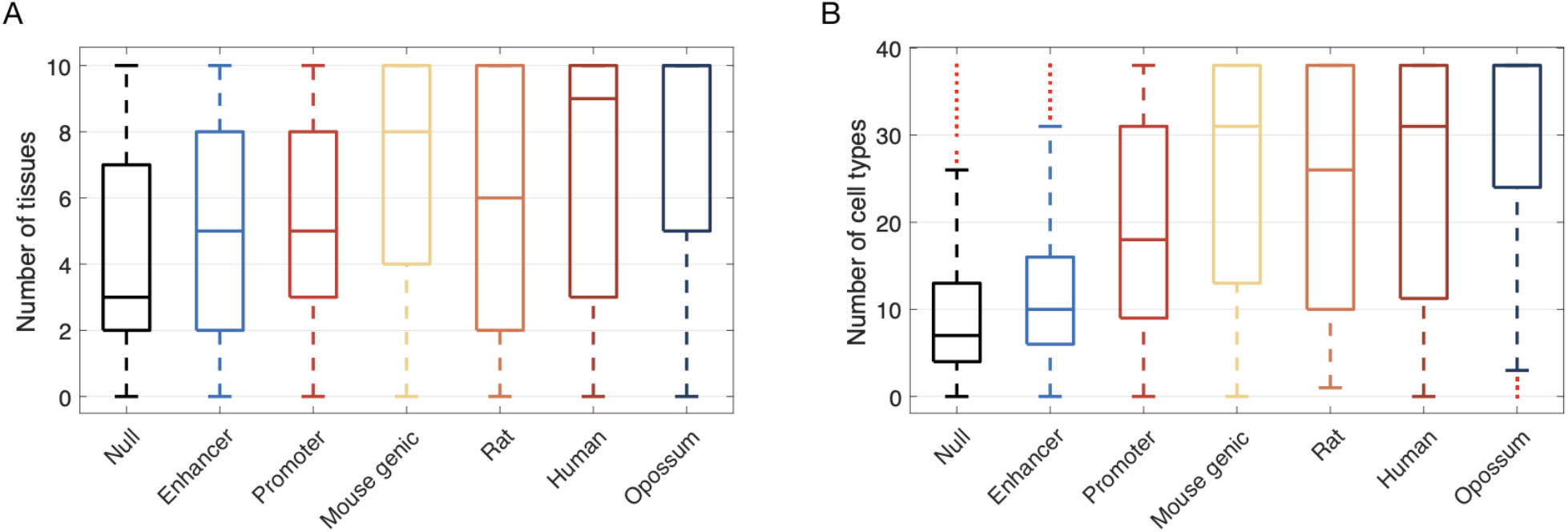
Mouse-specific intergenic ORFs that are proximal to enhancers are expressed in a limited diversity of cellular contexts. A) Number of tissues in which ORFs have an average FPKM > 0 across replicates. B) Number of cell types in which ORFs are in regions of open chromatin. In both panels, the “Null”, “Enhancer”, and “Promoter” categories correspond to mouse-specific intergenic ORFs.

### Some intergenic ORFs are proximal to promoters that show evidence of being repurposed enhancers

Similarities in the architectures of enhancers and promoters can facilitate the regulatory repurposing of enhancers into promoters (Wu and Sharp 2013; Carelli, et al. 2018), which could reinforce the transcription of ORFs emerging near enhancers. We next assessed whether the mouse-specific intergenic ORFs that are proximal to promoters are cases of ORFs transcribed from enhancers that were repurposed into promoters. To do so, we considered 329 mouse-specific intergenic ORFs that are expressed in mouse liver and proximal to an active promoter in mouse liver and that could be mapped to the rat genome via liftOver and (Fig. S2D; Materials and Methods). Subsequently, we assessed the chromatin modification status in the rat liver of those mapped genomic regions, using ChIP-seq data for H3K27ac and H3K4me3, marking enhancers and promoters, respectively.

Of the regions mapped to the rat genome, 256 are proximal to H3K27ac peaks and 183 are proximal to H3K4me3 peaks. The majority (70%) of the regions that are proximal to H3K27ac peaks are also proximal to H3K4me3 peaks (Fig. 5A,B), implying they act as promoters in the liver of both mouse and rat. However, some mapped genomic regions are at such distances from H3K4me3 peaks that they could well be enhancers in rat and may therefore have been repurposed into promoters on the lineage to mouse (Fig. 5B). Considering those mapped genomic regions with H3K27ac peaks that are separated from an H3K4me3 peak by a conservative threshold of at least 5kb, we found 33 candidates for the repurposing of rat enhancers to mouse promoters (∼10% of the 329 ORFs; Fig. 5B). The ORFs corresponding to these mapped genomic regions show evidence of stable transcription, both in terms of expression stability across biological replicates (Fig. 5C) and in terms of their proximity to capped analysis of gene expression (CAGE) peaks (Fig. 5D), which provides evidence that the transcripts are 5’-capped. Of note, many of these CAGE peaks are bidirectional, despite their proximity to H3K4me3 (promoter) peaks, which further supports the hypothesis of an enhancer origin (Fig. 5D). Finally, 79% of the 33 ORFs show evidence of association with ribosomes in mouse (using our most permissive criterion), which is more than we would expect by randomly sampling mouse-specific intergenic ORFs that are proximal to promoters and expressed in liver (Fig. 3C; binomial test *p* = 0.013). Together, these observations give further support to a model in which enhancers provide fertile ground for *de novo* gene birth.

**Figure 5.**
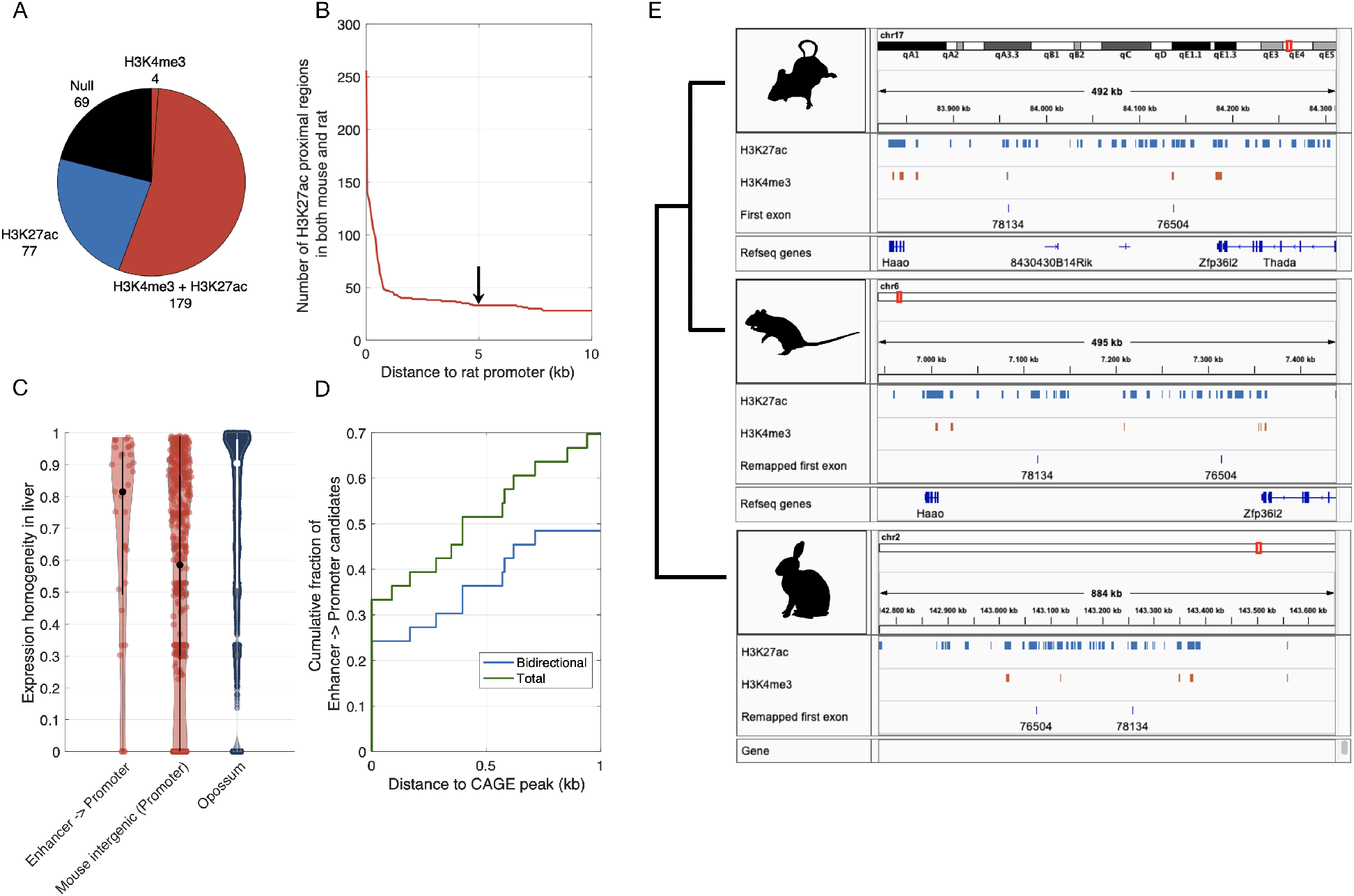
Some mouse-specific intergenic ORFs that are proximal to promoters show evidence of being repurposed enhancers. A) Distribution of histone methylation marks among genomic regions in rat. These regions are orthologous to genomic regions in mouse that harbour ORFs that are expressed in liver and are proximal to promoters. B) Number of genomic regions in rat liver with H3K27ac peaks, shown in relation to their distance to the closest H3K4me3 peak. We use a conservative threshold of 5kb (black arrow) between a promoter and an enhancer mark to determine that an enhancer is not a promoter. This results in 33 candidate promoters that were potentially repurposed from enhancers. C) Expression homogeneity in liver of these 33 ORFs (‘Enhancer -> Promoter’), mouse-specific intergenic ORFs that are proximal to promoters in liver, and opossum-shared ORFs. D) Cumulative fraction of the 33 ORFs shown in relation to their distance to the nearest CAGE peak (‘Total’) or the nearest bidirectional CAGE peak (‘Bidirectional’). E) Example repurposed enhancers. Orthologous genomic regions in mouse, rat, and rabbit that in mouse include the first exon of an intergenic mouse-specific ORF. The blue tracks represent H3K27ac peaks (enhancers) and the red tracks represent H3K4me3 peaks (promoters), both measured in liver samples from each organism. Annotated refseq genes are also indicated.

An alternative interpretation of these data is that promoters were repurposed as enhancers on the rat lineage, rather than enhancers being repurposed as promoters on the mouse lineage. To study the directionality of the repurposing, we considered ChIP-seq data for H3K27ac and H3K4me3 from the liver of rabbit, which served as an outgroup (Villar, et al. 2015). Of the 33 candidate genomic regions, 9 could be mapped to the rabbit genome using liftOver and were proximal to an H3K27ac peak in rabbit liver. Of these, 7 were proximal to an H3K27ac peak that was separated from an H3K4me3 peak by at least 5kb (see for example fig. 5E). This provides further support for the hypothesis of an ancestral enhancer state, at least for these 7 ORFs. Of note, all of the ORFs corresponding to these mapped genomic regions are surrounded by other enhancer marks in the mouse liver (Fig. S3), which hints that enhancer redundancy may help prevent conflicts that arise in the repurposing of enhancers into promoters, a possibility we revisit in the discussion.

### Enhancer interactions are gradually acquired over macro-evolutionary timescales

We next asked how enhancers integrate new genes into existing regulatory networks. To do so, we considered an enhancer-promoter interaction map derived from single-cell chromatin accessibility data in 13 murine tissues (Cusanovich, et al. 2018) (Materials and Methods). We uncovered a positive correlation between the age of an ORF and its number of enhancer interactions (Spearman’s correlation coefficient *ρ* = 0.23, *p* < 0.001; Fig. 6A). Among mouse-specific ORFs, intergenic ORFs had fewer enhancer interactions than genic ORFs, which were similar to non-mouse-specific ORFs in their number of enhancer interactions. This suggests that mouse-specific ORFs of genic origin tend to coopt the regulatory interactions of their host gene, or of nearby genes. Overall, the number of enhancer interactions increases from a median of 6 for mouse-specific intergenic ORFs to a median of 21 for ORFs that are shared with opossum, indicating that enhancer-promoter interactions are acquired gradually over time.

**Figure 6.**
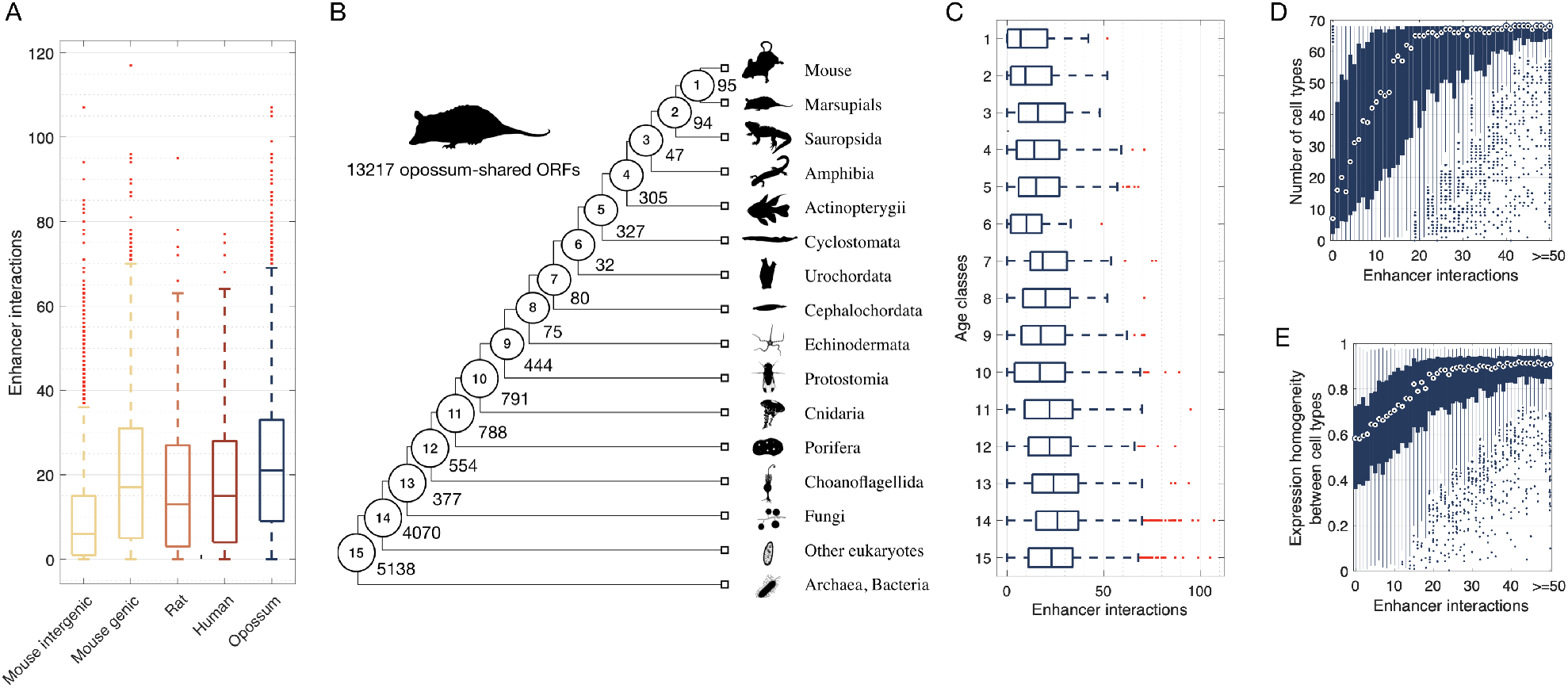
Enhancers facilitate the integration of genes into regulatory networks. A) Number of enhancer interactions per ORF. B) Phylogeny adapted from (Neme and Tautz 2013). The numbered circles indicate lineages representative of the age classes to which we assigned 13,217 opossum-shared ORFs, each of which corresponds to an annotated gene. The numbers on each branch represent the total number of annotated genes assigned to each age class. C) Number of enhancer interactions per gene, shown in relation to the age classes depicted in B. D) Expression breadth and E) homogeneity of opossum-shared annotated genes as a function of the number of enhancer interactions.

To explore the pace at which enhancer interactions are acquired over macro-evolutionary timescales, we shifted our focus to opossum-shared ORFs: we considered 13,217 opossum-shared ORFs corresponding to annotated genes with different first exons and separated them into 15 new age classes dating back to the origin of cellular life (Neme and Tautz 2013) (Fig. 6B). We again found a significant correlation between the age of a gene and its number of enhancer interactions (Spearman’s correlation coefficient *ρ* = 0.10, *p* < 0.001; Fig. 6C).

We next explored the consequences of enhancer acquisition. First, we studied the expression breadth of opossum-shared annotated genes using single-cell transcriptomic data from 68 cell types of ten murine tissues (Tabula Muris Consortium 2018), for which single-cell chromatin accessibility data were also available (Materials and Methods). We found that expression breadth increases with the number of enhancer interactions (Spearman’s correlation coefficient *ρ* = 0.49, *p* < 0.001; Fig. 6D) and with gene age (Spearman’s correlation coefficient *ρ* = 0.13, *p* < 0.001). The latter observation corroborates previous findings based on transcriptomic data from whole tissues (Kryuchkova-Mostacci and Robinson-Rechavi 2015). We next quantified the expression homogeneity of each gene across all cell types where expression was measurable (Materials and Methods), uncovering a positive correlation between expression homogeneity and the number of enhancer interactions (Spearman’s correlation coefficient *ρ* = 0.45, *p* < 0.001; Fig 6E), as well as gene age (Spearman’s correlation coefficient *ρ* = 0.11, *p* < 0.001).

Taken together, these results show that genes acquire enhancer interactions gradually over macro-evolutionary timescales, a process that correlates with expression breadth and homogeneity across cell types. Enhancers thus facilitate the integration of genes into regulatory networks.

## Discussion

Our study provides empirical support for the hypothesis that enhancers may facilitate *de novo* gene evolution, which to our knowledge was first proposed upon the discovery of enhancer RNA (Kim, et al. 2010) and later expanded upon in a perspective piece by Wu and Sharp (Wu and Sharp 2013). Our findings complement recent work on the regulatory architecture of the nematode *Pristionchus pacificus,* which showed that young genes – those private to *P. pacificus* – are in closer proximity to enhancers than genes with one-to-one orthologs in other nematode species (Werner, et al. 2018). The observation that many young ORFs are proximal to enhancers in both nematodes and mammals suggests that this mode of gene evolution dates back to at least the common ancestor of Bilateria, and possibly even earlier, since cnidarians, ctenophores, and sponges also employ distal regulatory elements (Schwaiger, et al. 2014; Gaiti, et al. 2017; Sebé-Pedrós, et al. 2018; Sebé-Pedrós, et al. 2018). As the complexity of gene regulation increased during the evolution of some lineages, such as the lineage to vertebrates (Marletaz, et al. 2018), we speculate that enhancer-facilitated *de novo* gene birth may have played an increasingly prominent role in the expansion of gene regulatory networks.

The facilitating role of enhancers in *de novo* gene birth is conceptually similar to the facilitating role of the permissive chromatin state of meiotic spermatocytes and post-meiotic round spermatids that underlies the “out-of-testis hypothesis,” which proposes the testis as a primary tissue for the origination of new genes (Kaessmann 2010; Witt, et al. 2019). Both scenarios envision regions of open chromatin that are exposed to the transcriptional machinery, and thus produce a transcriptionally active environment that is conducive to the evolution of new genes. The two scenarios differ, however, in at least three ways. First, genes that emerge from or near enhancers may rapidly acquire their own promoters, due to the similar architectural and functional features of enhancers and promoters, a similarity that facilitates the repurposing of the former to the latter (Wu and Sharp 2013). Indeed, we report several mouse-specific intergenic ORFs that are proximal to promoters that show evidence of being repurposed enhancers, complementing recent analyses of enhancer repurposing in primates and rodents (Carelli, et al. 2018). Second, enhancers are often deployed in multiple cell types or developmental stages (Kvon, et al. 2014), exposing enhancer-proximal young ORFs to selection in a limited diversity of cellular contexts. This may help to purge toxic peptides (Wilson and Masel 2011), and balance the benefit of expression in distinct cellular environments with the cost of pleiotropic effects. Third, because enhancers are often active in somatic cell types, *de novo* genes emerging near enhancers are more likely to be involved in physiological or morphological traits than *de novo* genes emerging from testis, which are more likely to be involved in reproductive traits. We emphasize that the enhancer-facilitated and out-of-testis scenarios are not mutually exclusive; in fact, they may be complementary or even interactive. Indeed, we found many young transcribed ORFs that associate with ribosomes in testis that are also proximal to enhancers (Figs. S2F, 3C).

The three points that differentiate enhancer-facilitated *de novo* gene birth from the out-of-testis scenario also differentiate enhancer-facilitated *de novo* gene birth from pervasive transcription (Clark, et al. 2011) taking place away from promoters or enhancers. An additional difference is the relatively high and stable expression levels of enhancers, which increases the chances of ORF-bearing transcripts that stem from or near enhancers to associate with ribosomes. This is indeed what we observe when comparing ribosomal association among mouse-specific intergenic ORFs that are proximal to enhancers with mouse-specific intergenic ORFs that are not proximal to enhancers or promoters. However, we note that this observation may be a technological artifact. If the likelihood of the ribo-seq assay to detect ribosomal association increases with the level or stability of expression, then we would expect to see increased ribosomal association for mouse-specific intergenic ORFs that are proximal to enhancers, relative to mouse-specific intergenic ORFs that are not proximal to enhancers or promoters, even if these two classes of ORFs tend to associate with ribosomes to the same extent. Thus, the reason why enhancer proximity increases the likelihood of ribosomal association is the same reason why we cannot rule out the possibility that we observe this association due to a technological artifact.

An additional facet to enhancer-facilitated *de novo* gene birth is conflict between the enhancer and the emerging gene. If the enhancer is repurposed as a promoter to enforce directional transcription, then the ancestral function of the enhancer may be compromised. There are at least two ways to resolve this conflict. One is to maintain enhancer function; indeed, many promoters also act as enhancers (Medina-Rivera, et al. 2018). Another is enhancer redundancy. Genes are often targeted by multiple enhancers, and in many of these cases, only a subset of the enhancers are necessary to drive correct expression under normal growth conditions (Osterwalder, et al. 2018). Thus, we hypothesize that redundant enhancers are less likely to face conflict in facilitating *de novo* gene birth. While our observation that repurposed enhancers tend to be surrounded by other enhancers provides anecdotal support for this hypothesis (Fig. S3), more systematic analyses are warranted.

The hypothesis that enhancers help *de novo* genes integrate into existing regulatory networks was previously proposed in the context of the out-of-testis hypothesis, as a means to expand a new gene’s breadth of expression (Tautz and Domazet-Loso 2011). Using single-cell chromatin accessibility and transcriptomic data, our study provides empirical support for the hypothesis that *de novo* genes gradually acquire enhancer interactions over time, and that this acquisition increases expression breadth and homogeneity. These findings complement related studies of gene integration into cellular networks, such as networks of protein-protein interactions (Capra, et al. 2010; Abrusán 2013). Our observation that genes continue to acquire enhancer interactions over macro-evolutionary timescales mirrors similar increases in other aspects of gene regulation, such as in the number of proximal transcription factor binding sites, alternative transcript isoforms, and miRNA targets (Warnefors and Eyre-Walker 2011).

Regulatory networks drive the spatiotemporal gene expression patterns that give rise to and define the numerous and distinct cellular identities characteristic of Metazoan life. Enhancers play an integral role in this process, mediating cell-type-specific gene-gene interactions, thus facilitating the combinatorial deployment of different genes and network modules in different contexts. Genetic changes that affect such interactions are responsible for myriad evolutionary adaptations and innovations (Carroll 2001; Prud’homme, et al. 2007; Carroll 2008; Peter and Davidson 2011). Our results suggest that the power of enhancers in creating such evolutionary novelties lies not only in their ability to rewire gene regulatory networks, but also in their ability to expand them, by providing fertile ground for *de novo* gene birth.

## Materials and methods

### ORF age and classification

Schmitz et al. (2018) identified a set of 58,864 ORFs from the transcriptomes of three murine tissues: liver, brain, and testis. Blasting against the transcriptomes of four other mammalian species (rat, human, kangaroo rat, and opossum), they estimated the age of each ORF by phylostratigraphic methods (Domazet-Lošo, et al. 2007; Schmitz, et al. 2018). Because of the small number of ORFs shared with the kangaroo rat (49 ORFs), we merged these ORFs together with those from the rat age class. We used the genomic coordinates of the first exon of each ORF in the mm10 mouse genome reference to study the regulatory properties of ORFs of different ages, for example to study their distance to the nearest enhancer. We only considered ORFs that were transcribed from nuclear chromosomes and whose first exon was longer than 30 base pairs. If first exons were shared between more than one ORF, we only retained the oldest of the ORFs. Our filtered dataset contained 56,262 ORFs.

Schmitz et al. (2018) annotated each ORF as belonging to one of 8 different categories: “intergenic,” “close to promoter same strand,” “close to promoter opposite strand,” “overlapping same strand,” “overlapping opposite strand,” “overlapping coding sequence same strand,” “overlapping coding sequence opposite strand,” and “overlapping annotated gene in frame.” We considered all categories except “intergenic” to be “genic” in order to separate ORFs that were born within or near existing genes from those that were not. This resulted in 5 classes: Mouse-specific intergenic ORFs, mouse-specific genic ORFs, rat-shared ORFs, human-shared ORFs and opossum-shared ORFs.

### Proximity to enhancers and promoters

We obtained ChIP-seq data for H3K27ac, H3K4me1, and H3K4me3 modifications from 23 different tissues and cell types from the ENCODE project (bone marrow, cerebellum, cortex, heart, kidney, liver, lung, olfactory bulb, placenta, spleen, small intestine, testis, thymus, embryonic whole brain, embryonic liver, embryonic limb, brown adipose tissue, macrophages, MEL, MEF, mESC, CH12 cell line, and E14 embryonic mouse) (ENCODE 2012). We used liftOver (Kent, et al. 2002) to convert the genomic coordinates of the peaks from mm9 to mm10. We used the “merge” function of bedtools (Quinlan and Hall 2010) with default parameters to collate the peaks for all tissues and cell types, considering any overlapping H3K27ac and H3K4me1 peak as part of the same enhancer. We used the “intersect” function of bedtools with default parameters to separate H3K27ac and H3K4me1 peaks that overlapped any length of H3K4me3 peaks from those that did not. This resulted in 172,930 H3K27ac and 277,187 H3K4me1 peaks that did not overlap H3K4me3 peaks. We considered genomic regions with H3K4me3 peaks to be promoters, and those exclusively with H3K27ac and/or H3K4me1 peaks to be enhancers (Berthelot, et al. 2018). We measured the distance in base pairs between the first exon of an ORF to an enhancer or promoter using the “closest” function of bedtools with the “-t first” option activated. We considered an ORF to be proximal to an enhancer if the distance to the first exon was shorter than 500 bp and there was no promoter within that distance. When controlling for the length of the first exon, we considered the distance to windows of 750 bp up-and downstream of the central nucleotide of the first exon, rather than to the first exon itself.

We followed the same procedures when measuring the distance of ORFs to enhancers and promoters in liver, brain, and testis tissues separately. For brain tissue, we merged ChIP-seq data from embryonic whole brain and cortex. The ORFs we considered as expressed in each tissue (Fig. S2A-C) were those with a mean FPKM greater than zero across replicates of total RNA transcriptomic data (8 replicates for liver and brain, and 2 replicates for testis) (Li, et al. 2017).

### Chromatin accessibility

We used single-cell ATAC-seq data from 13 different mouse tissues (bone marrow, cerebellum, large intestine, heart, small intestine, kidney, liver, lung, cortex, spleen, testes, thymus, and whole brain). We obtained the data from the Mouse ATAC atlas (Cusanovich, et al. 2018), which comprised 436,206 peaks of open chromatin. We used liftOver to convert the genome coordinates from mm9 to mm10. A total of 29 peaks could not be converted. Using the “closest” function of bedtools with the “t first” option activated, we calculated the distance between ORFs and regions of open chromatin. We annotated regions of open chromatin as enhancers if they overlapped H3K27ac and/or H3K4me1 peaks but not H3K4me3 peaks, or as promoters if they overlapped H3K4me3 peaks. To do so, we used the bedtools intersect function with the -u option activated.

Cusanovich et al. (2018) used these single-cell ATAC-seq data to identify clusters of cells with similar patterns of chromatin accessibility. They assigned the clusters to 38 distinct cell types based on the chromatin accessibility of marker genes indicative of each cell type. We used these data to identify tissues and cell types where ORFs are in accessible chromatin. We considered an ORF-containing region of the genome to be in open chromatin in a certain cell type if it was accessible in at least 1% of the cells that made up at least one of the clusters of that cell type (Cusanovich et al. 2018).

### 5’capping

We used cap analysis of gene expression (CAGE) data from the FANTOM5 consortium from 1,016 mouse samples including cell lines, primary cells, and tissues (Lizio, et al. 2015; Noguchi, et al. 2017). This method is based on the capture of 5’ capped ends of mRNA, which allows the mapping of regions of transcription initiation genome-wide (Shiraki, et al. 2003). Using the “closest” function from bedtools with the “-t first” option activated (Quinlan and Hall 2010), we measured the distance between an ORF’s first exon and its closest CAGE peak. In the same manner, we also considered a subset of CAGE peaks which were annotated as bidirectional and transcribed from enhancers (Andersson, et al. 2014; Dalby, et al. 2018).

### Expression level and stability

We measured the expression levels and stabilities of ORFs. To do so, we aligned paired reads produced by RNAseq from total RNA from 10 tissues (liver, testis, brain, muscle, bone, small intestine, thymus, heart, lung and spleen) (Li, et al. 2017) using STAR 2.5.3a (Dobin, et al. 2013) to the mm10 build of the mouse genome. We chose these tissues because ChIP-seq data for histone modifications were also available. For each ORF, we calculated fragments per kilobase per million mapped reads (FPKM) as the number of reads mapped to the first exon divided by a millionth of the number of reads sequenced in each sample and then by the length of the exon in kilobases. We considered an ORF to be expressed if it had an average FPKM greater than 0 across replicates.

We calculated the expression stability of ORF *k* as:

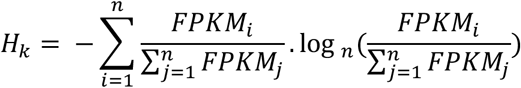

where *n* is the number of replicates for a given tissue (8 for liver and brain, 2 for testis). We refer to this measure as expression homogeneity when calculated across tissues or cell types, rather than across replicates for the same tissue or cell type.

### Ribosome association

We used ribosome profiling (ribo-seq) data from mouse liver, brain, and testis (Ingolia 2014). We obtained the coordinates of mRNA segments detected by ribo-seq from GWIPS-viz (Michel, et al. 2014), a database that includes such data from different studies. From this source, we considered samples from liver (3 samples from 3 studies), brain (5 samples from 2 studies), and testis (1 sample) (details provided in Supplementary text 1). We combined the datasets for each tissue and merged the provided genomic coordinates using the bedtools merge function; we did so with the options “-c” and “-o absmax” activated. Following Ruiz-Orera et al. (2018), we removed all merged coordinates shorter than 26 bp, because these could be anomalous reads. We subsequently mapped these merged coordinates on the first exon of our set of ORFs using the bedtools “map” function and we summed the number of reads from each of the mapped merged coordinates. In this way, we were able to assign a number of ribo-seq reads to each ORF, which allowed us to estimate ribosomal association and thus potential for translation.

### Enhancer repurposing

We considered the set of 544 mouse-specific intergenic ORFs that were transcribed (average FPKM > 0) and proximal to an H3K4me3 peak in mouse liver. We filtered this set to the 357 ORFs that were proximal to what Villar et al. (2015) considered to be replicated H3K4me3 peaks in mouse, in order to facilitate comparison with the histone methylation data from rat and rabbit that were generated for the same study. We used liftOver to map the genomic coordinates of these ORFs to the rat and rabbit genomes (builds r5 and oryCun2), requiring a minimum fraction of remapped bases of 0.6 and 0.4 respectively (Carelli, et al. 2018). This resulted in 329 and 130 presumably orthologous genomic regions in rat and rabbit respectively. Considering H3K27ac and H3K4me3 ChIP-seq peaks in the livers of mouse, rat, and rabbit (Villar, et al. 2015), we then calculated the distance between these mapped regions and H3K4me3 and H3K27ac peaks using the bedtools “closest” function. We considered the promoter of an ORF in mouse to show evidence of being a repurposed enhancer if its mapped genomic region in rat or rabbit was proximal to an H3K27ac peak, yet more than 5 kb from an H3K4me3 peak, in rat or rabbit liver.

### Enhancer interactions

Cusanovich et al. (Cusanovich, et al. 2018) used single-cell ATAC-seq data to predict physical interactions between regions of open chromatin (Pliner, et al. 2018), thus creating an atlas of enhancer interactions in single murine cells. We downloaded these data from the Mouse ATAC atlas (Cusanovich, et al. 2018), which includes the cell clusters where the interactions occur, as well as the co-accessibility scores of pairs of regions of open chromatin – a measure of interaction strength. We disregarded cell clusters classified as “unknown” or “collisions”, as well as interactions with a co-accessibility score lower than 0.25, following Pliner et al. (2018). We also filtered out interactions with regions of open chromatin that overlapped ChIP-seq peaks for H3K4me3 marks or no enhancer marks, in order to focus solely on interactions with enhancers. An interaction was assigned to an ORF if the ORF’s first exon was included in the interaction.

### Age of annotated genes

To study how genes acquire enhancer interactions over macro-evolutionary timescales, we considered the subset of ORFs that belong to the opossum age class in Schmitz et al. (2018) and that are annotated as genes in the latest version of Ensembl (release 95) (Cunningham, et al. 2019). We matched these genes to age estimates reported by Neme & Tautz (2013), based on a phylostratigraphic analysis of 20 lineages spanning 4 billion years from the last universal common ancestor to the common ancestor of mouse and rat. We further filtered the dataset to only include ORFs that emerged in the first 15 of the 20 phylostrata, in order to focus on ORFs that are considered to have emerged before the split between the common ancestor of placental mammals and marsupials by both Schmitz et al. (2018) and Neme & Tautz (2013). This left us with ∼16,000 ORFs corresponding to 13,217 unique annotated genes that emerged prior to the origin of placental mammals.

### Expression breadth and homogeneity of annotated genes

To study the transcription of annotated genes, we used the expression data reported by the Tabula Muris Consortium (Tabula Muris Consortium 2018) for the single-cell RNA sequencing performed with FACS-based cell capture in plates, for 20 different mouse tissues. The data include the log-normalization of 1 + counts per million for each of the annotated genes in each of the sequenced cells. We considered ten tissues that were also used for the construction of the Mouse ATAC Atlas (Cusanovich, et al. 2018). We measured the expression breadth of each ORF corresponding to an annotated gene as the number of cell types in which expression could be detected in at least 1% of the cells assigned to a cell type. The homogeneity of expression across cell types was calculated as explained above for *Hk*, but considering the mean expression in each cell type where expression was detectable, rather than the mean expression across replicates.

## Supplementary figures

**Figure S1.**
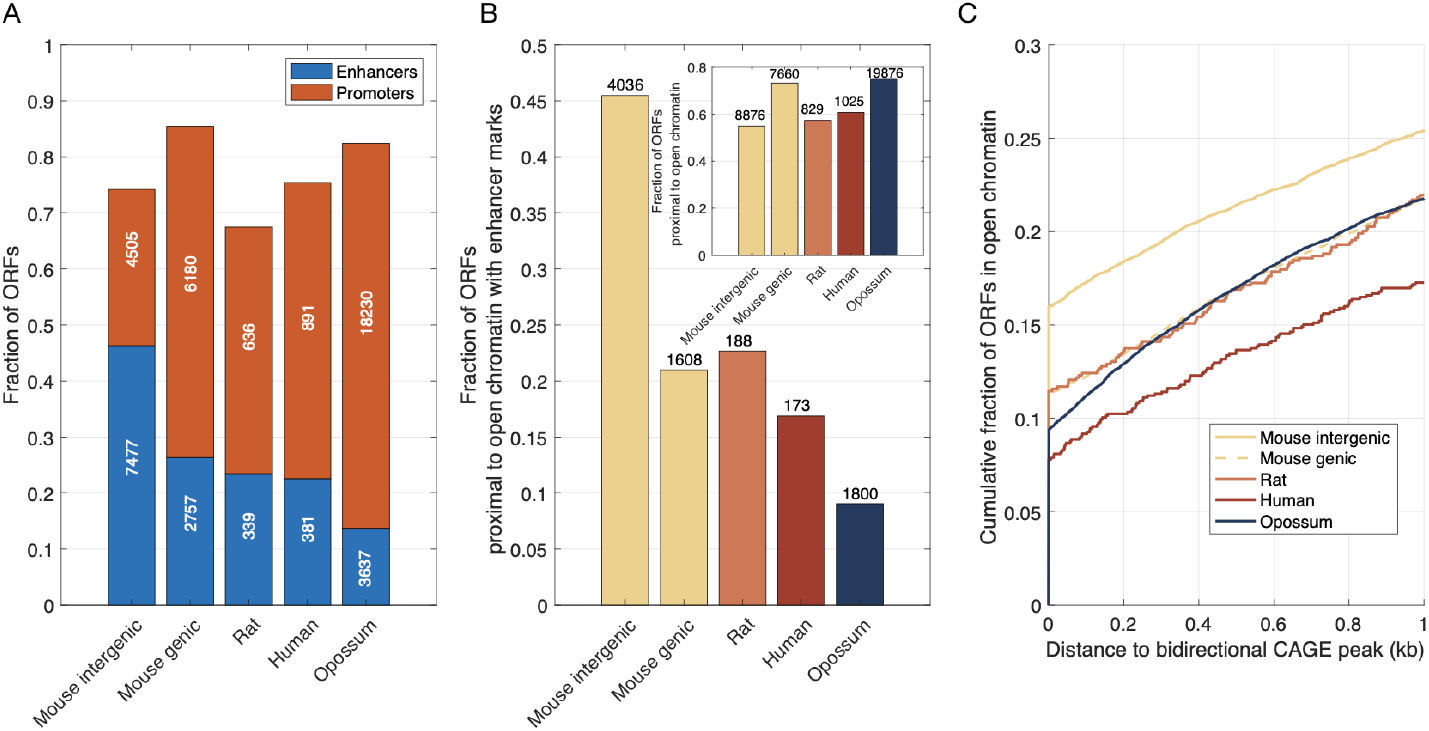
Exon length is not responsible for the proximity of young ORFs to enhancers. Repeating the analyses shown in Fig. 1B-D using windows of 1.5 kb around the central nucleotide of the first exon of each ORF, rather than the first nucleotide of the first exon, results in the same qualitative trend that mouse-specific intergenic ORFs are more likely to be proximal to enhancers than the other classes of ORFs studied here.

**Figure S2.**
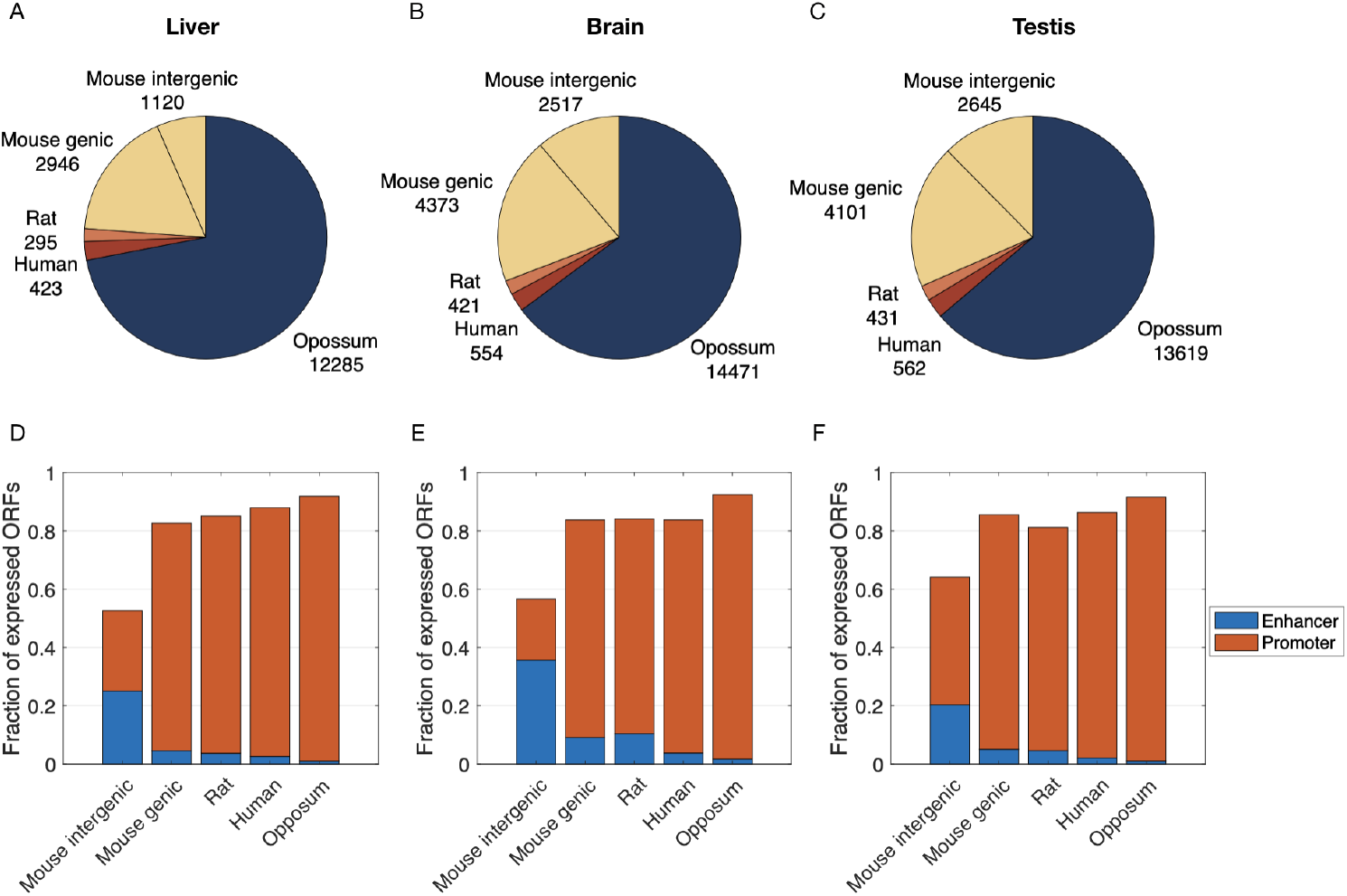
Mouse-specific intergenic ORFs are more associated with active enhancers in individual tissues than older and genic ORFs. A-C) Number of ORFs of each class that are in regions of open chromatin and that are expressed (FPKM > 0) in (A) liver, (B) brain, and (C) testis. D-F) Fraction of ORFs from each class (as presented in A-C) that are proximal to promoters or enhancers in (D) liver, (E) brain, and (F) testis.

**Figure S3.**
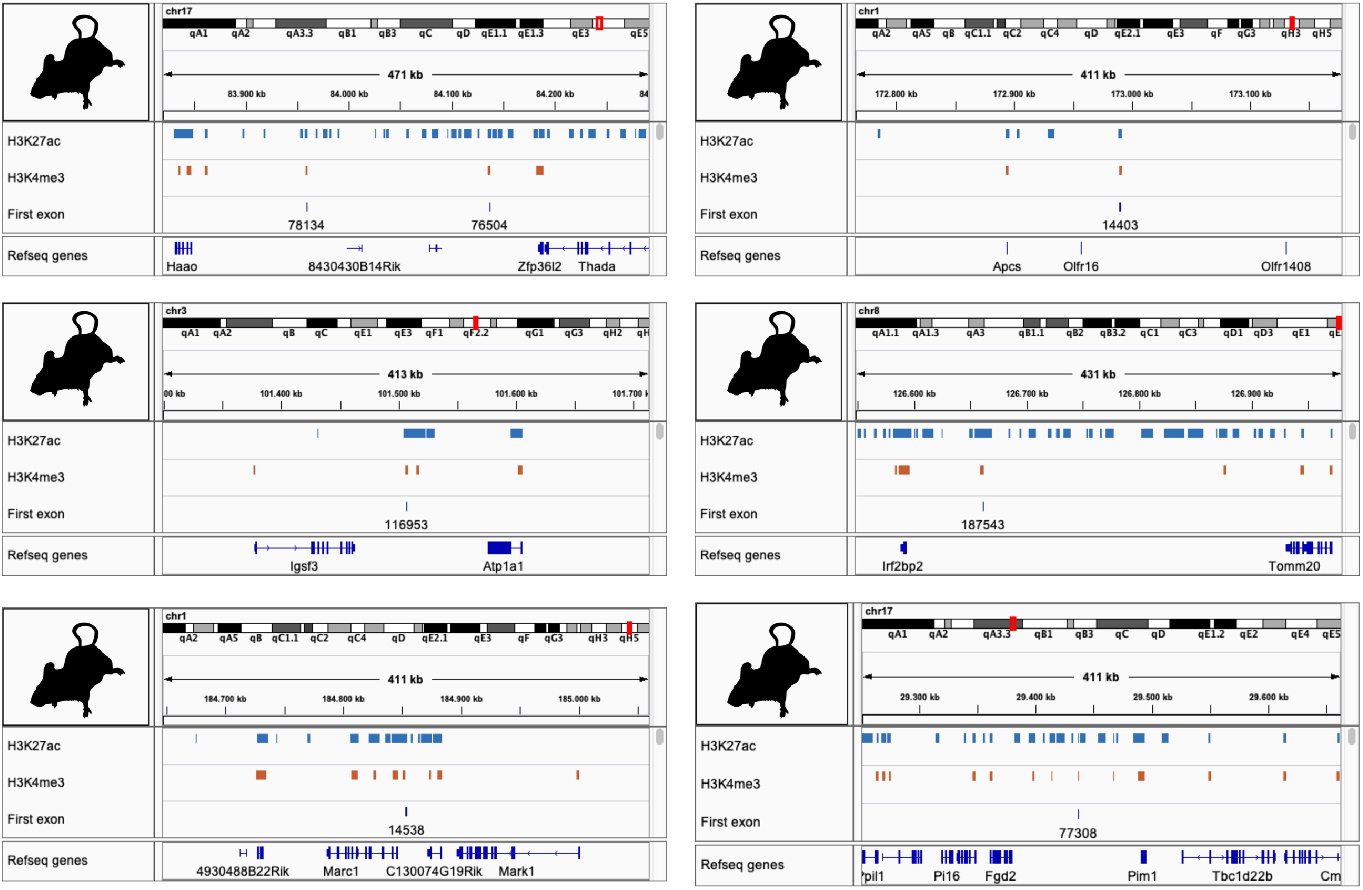
Enhancer redundancy may help prevent conflicts in the repurposing of enhancers to promoters. Visualization of 6 genomic regions that contain 7 mouse-specific intergenic ORFs that are proximal to promoters and show evidence of being repurposed enhancers.

**Figure S4.**
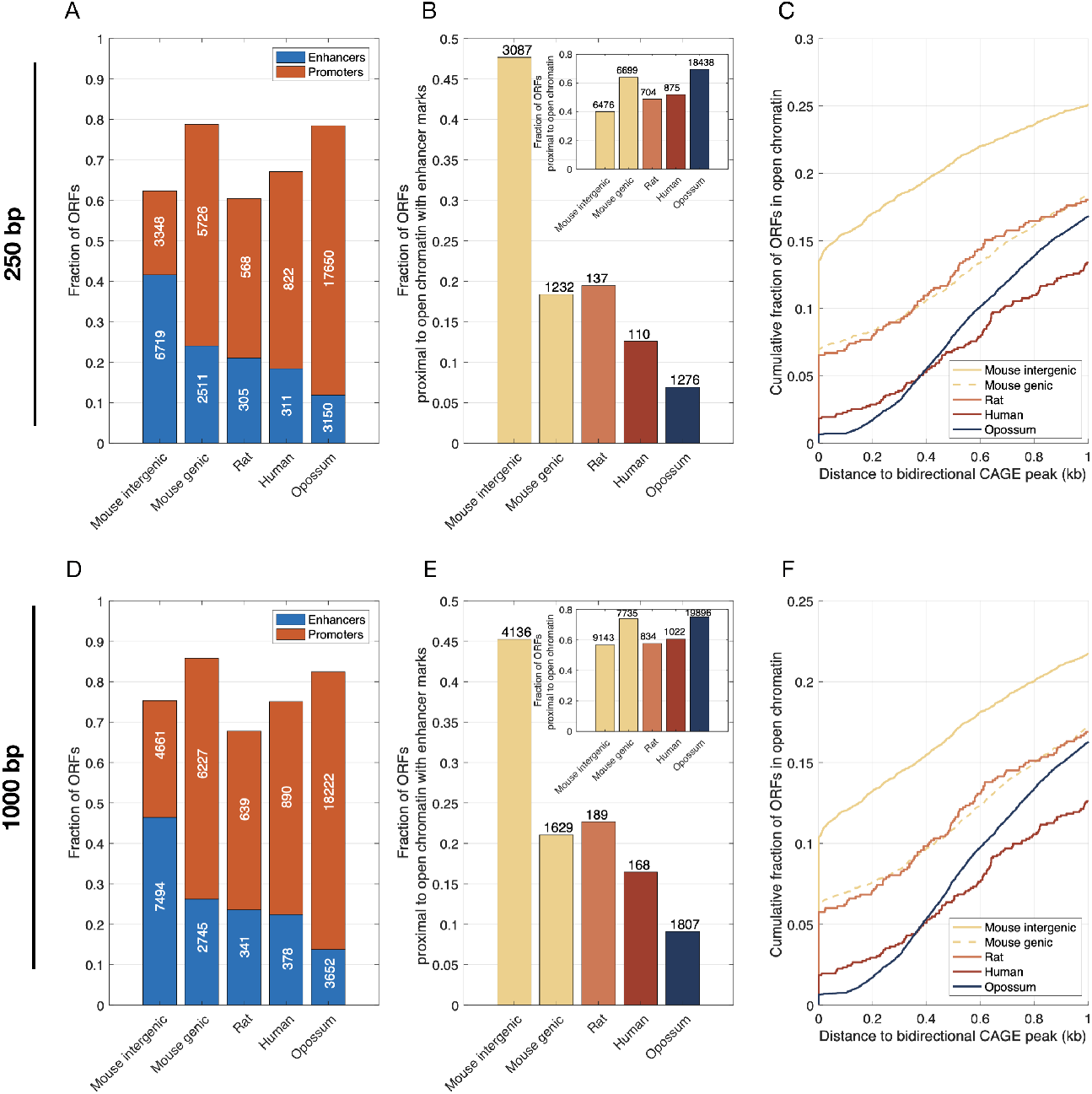
Changing our definition of proximal from within 500bp to within (A-C) 250bp or (D-F) 1000bp does not qualitatively alter the trends shown in Fig. 1B-D.

**Figure S5.**
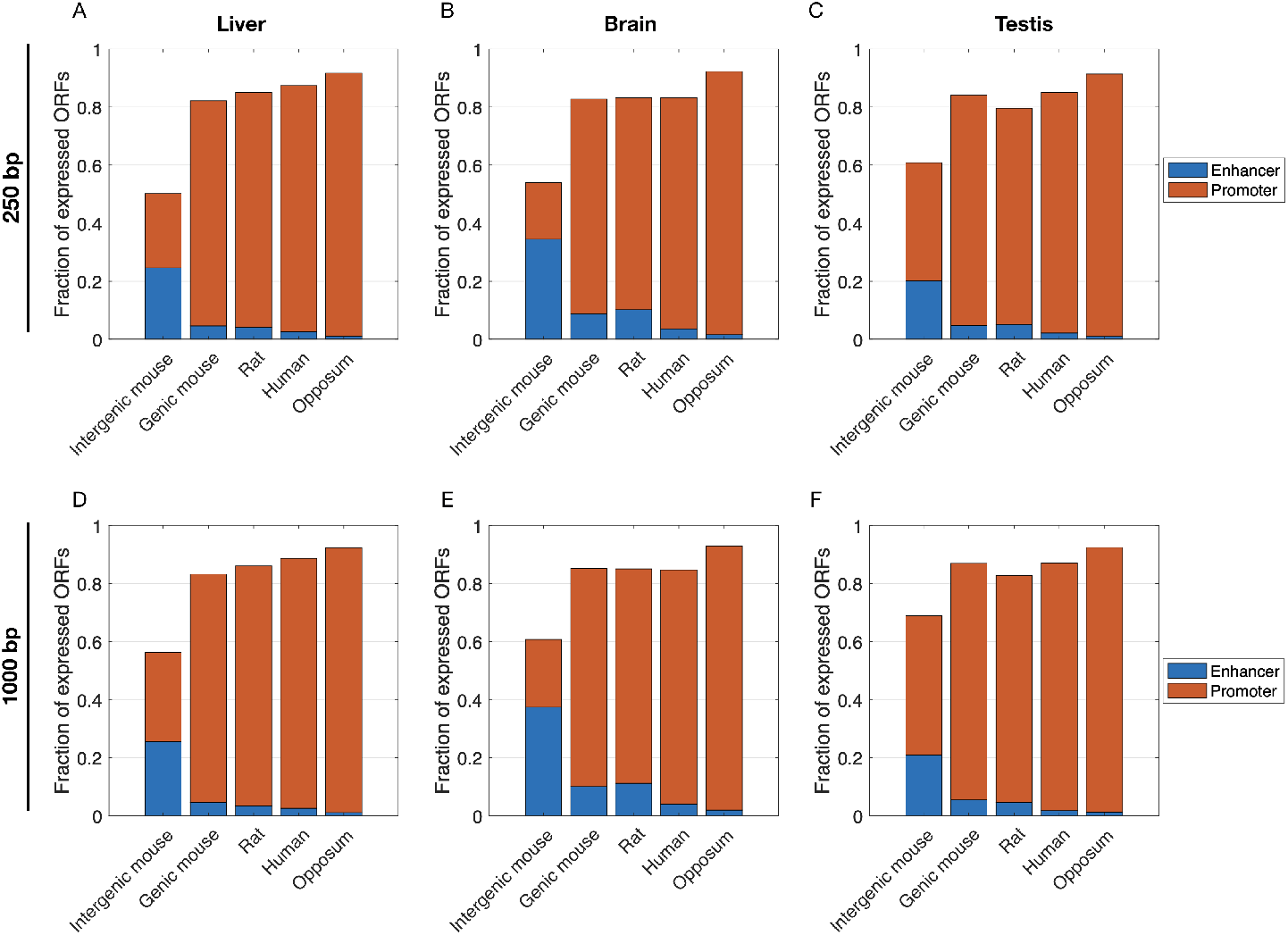
Changing our definition of proximal from within 500bp to within (A-C) 250bp or (D-F) 1000bp does not qualitatively alter the trends shown in Fig. S2D-F.

**Figure S6.**
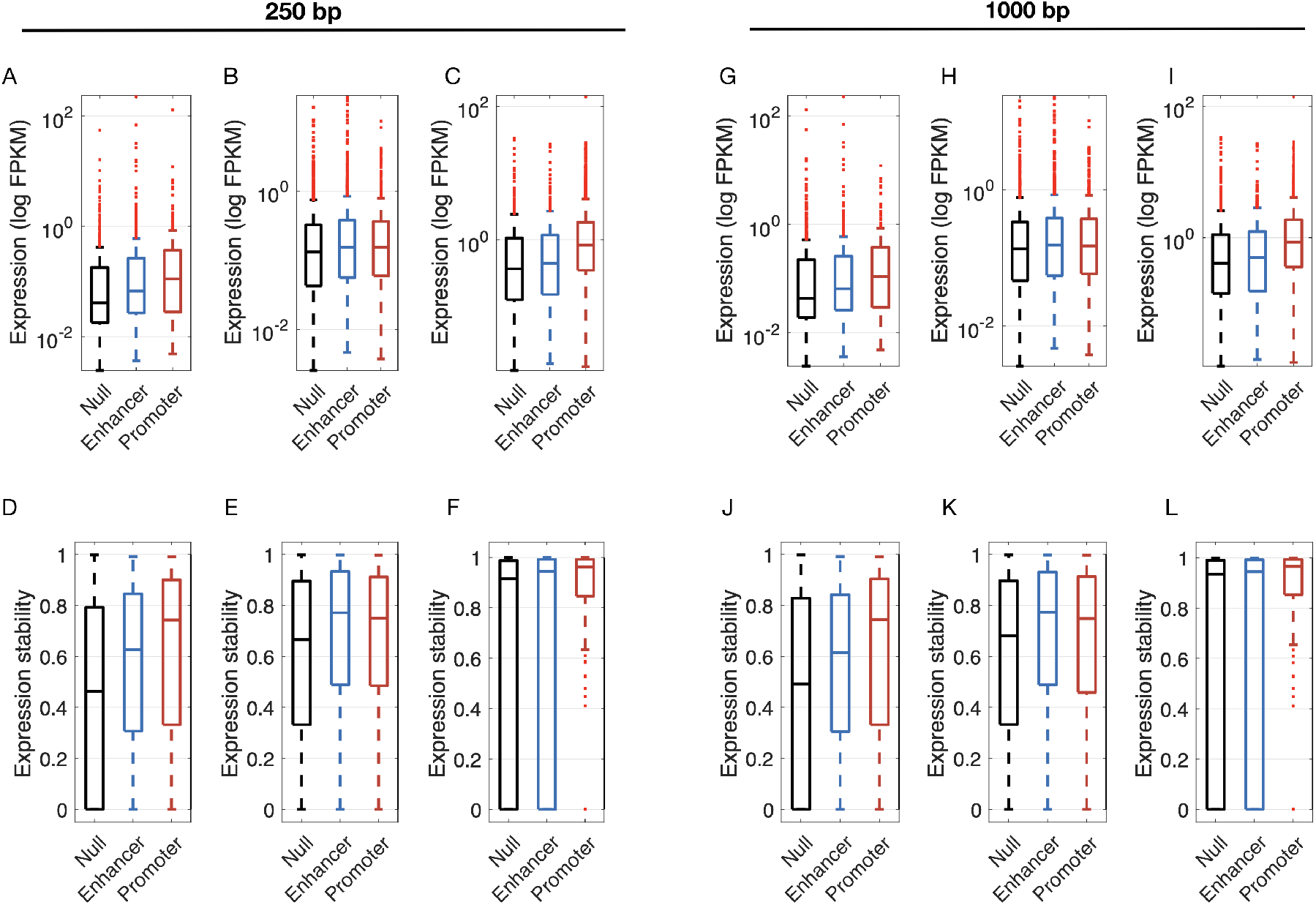
Changing our definition of proximal from within 500bp to within (A-F) 250bp or (G-L) 1000bp does not qualitatively alter the trends shown in Fig. 2.

**Figure S7.**
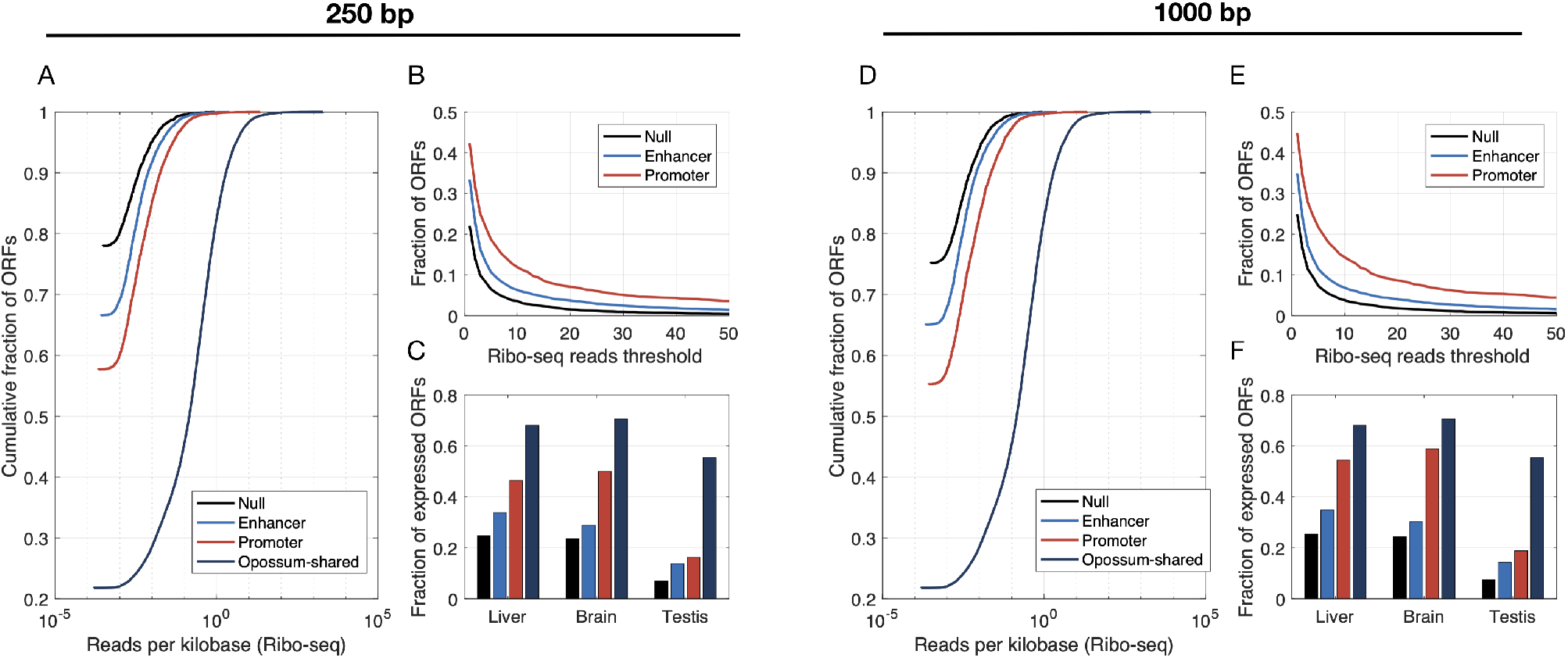
Changing our definition of proximal from within 500bp to within (A-C) 250bp or (D-F) 1000bp does not qualitatively alter the trends shown in Fig. 3.

**Figure S8.**
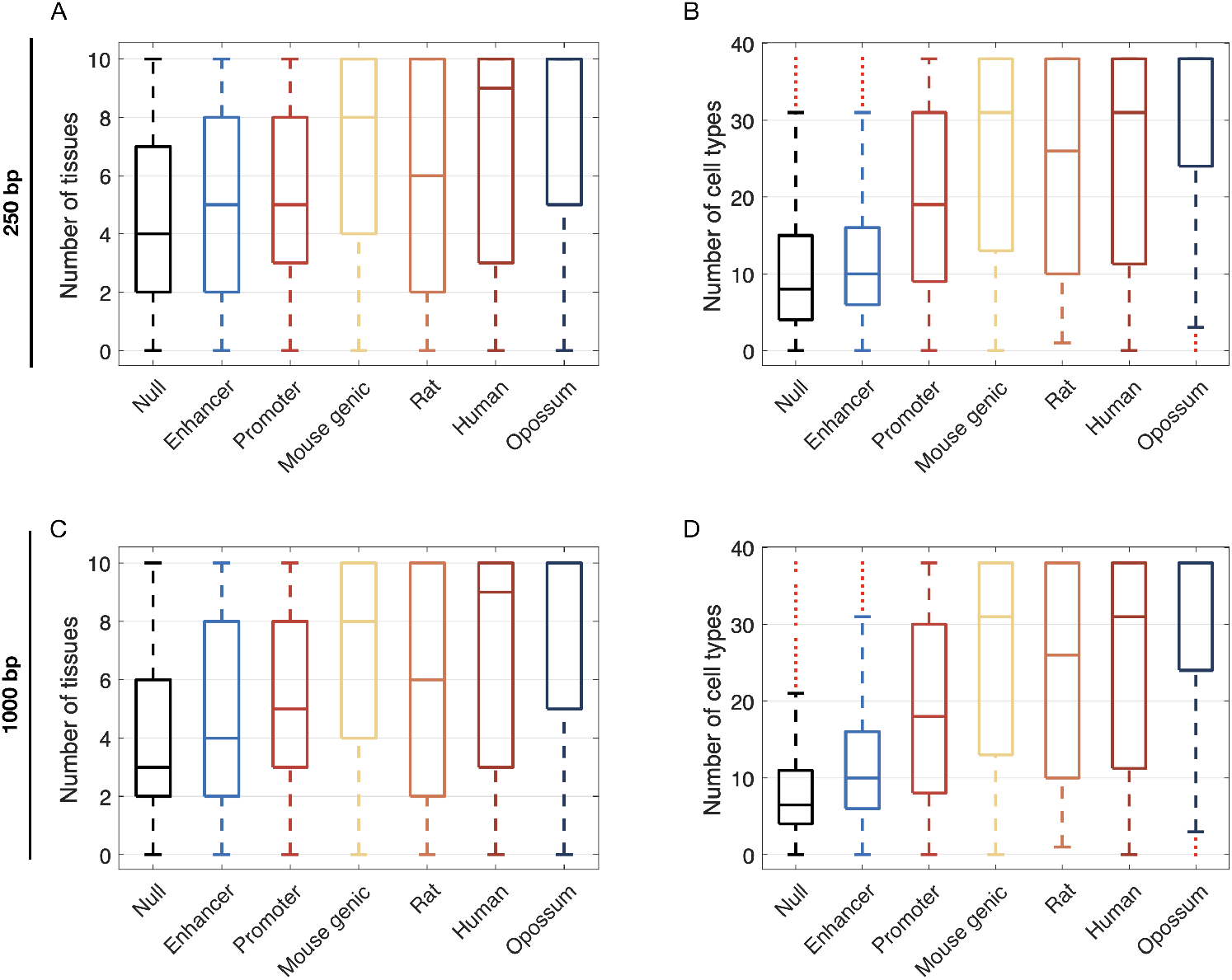
Changing our definition of proximal from within 500bp to within (A,B) 250bp or (C,D) 1000bp does not qualitatively alter the trends shown in Fig. 4.

